# Intersection of genotype and environment on the dynamics of the virome and microbiome of an estuarine cnidarian

**DOI:** 10.64898/2026.07.23.740364

**Authors:** Sydney Birch, Yehu Moran, Adam M. Reitzel

## Abstract

Estuaries host a rich diversity of microorganisms, creating opportunities for animal–microbe interactions that shape host physiology. The sea anemone *Nematostella vectensis* spans the eastern coast of North America from Nova Scotia to Florida, and prior work shows that anemones from different regions harbor distinct microbial communities. Although the virome of the standard laboratory population has been characterized, population-level variation in viromes remains largely unknown. To address this gap, we conducted a mesocosm experiment using six *N. vectensis* populations to test whether geographic origin influences the diversity and functional potential of associated viruses and bacteria. We sampled individuals before mesocosm exposure and 14 days after to sequence their viromes, microbiomes, and host transcriptomes. We also collected anemones from a natural population to compare against experimental animals. Both virome and microbiome analyses revealed increased taxonomic and functional diversity after 14 days, with the strongest increase observed in lower-latitude populations. Notably, experimental anemones did converge toward the natural population’s community composition; however, population-specific interactions with viruses and microbes persisted. Differential gene expression indicated a modest host response overall, whereas WGCNA identified a clear north–south expression pattern. Together, these findings demonstrate that *N. vectensis* genotypes from locations along a latitudinal gradient maintain distinct viral and microbial associations, highlighting geographically structured host–microbe–virus relationships.

## Introduction

Marine environments are home to a diverse array of microorganisms, including bacteria, archaea, viruses, and fungi, enabling animal-microbe interactions. Many investigations have focused on the bacterial-animal associations across various marine invertebrate taxa, shedding light on the importance of microbes on host physiology ^1–4^. However, less is known about how assemblages of viruses (viromes) colonize and impact organisms, particularly in marine and aquatic species, outside aquaculture or those known to commonly vector pathogenic viruses to humans ^5^.

As the most abundant and diverse biological entities on Earth ^5–8^, viruses can exert both direct and indirect effects on host animals. Direct effects arise from infection, which may cause disease or mortality, whereas indirect effects can occur through virus-mediated alterations of the host-associated microbiome, with downstream consequences for host physiology ^9–12^. Advances in metagenomic and metatranscriptomic sequencing have greatly expanded our understanding of viral diversity associated with non-arthropod marine invertebrates, which have historically received disproportionate attention compared to arthropod invertebrates due to their role as vectors of human pathogens ^13,14^. These studies reveal that marine invertebrate viromes are often more diverse than those of vertebrates^15,16^, raising fundamental questions about the functional roles of these viruses, their impacts on host biology, and the extent to which virome composition varies among populations within a single species.

The phylum Cnidaria, which encompasses sea anemones, corals, jellyfish, and hydrozoans, provides a powerful marine system for investigating interactions among microbiomes, viromes, and host animals. Cnidarians include ecologically important species, such as reef-building corals, that are increasingly impacted by viral and microbial diseases ^17^. Additionally, many anthozoans (corals and sea anemones) harbor complex microbial communities composed of bacteria, unicellular eukaryotes, and fungi, a feature that has motivated extensive research in recent years ^18^. Recent studies have begun to identify crosstalk between eukaryotic and prokaryotic microbiota, particularly for reef-building corals ^19^. Additionally, bacterial microbiomes in anthozoans have been shown to play a role in nutrient cycling ^20–24^, protecting against pathogens ^25–31^, influencing metabolic processes ^32–34^, aiding in response to environmental fluctuations ^35^, and regeneration ^36^.

Previous research has examined how cnidarian microbiomes vary across populations and closely related species ^18^. Studies of sea anemones have shown that individuals collected from different geographic locations possess distinct microbiomes (e.g., *Nematostella vectensis* ^37^, *Anthopleura elegantissima* ^38^, but see ^39^). Similarly, acroporid corals exhibit distinct microbiomes that vary by location but not between species ^40^, suggesting that location may play a major role in microbiome composition. This has also been observed in other reef-building corals ^41,42^, further supporting that bacterial diversity correlates with location and environmental conditions. Genetically distinct strains within a species have also been shown to harbor unique bacterial communities^43^, and recent work suggests that genetic variation among coral populations contributes to adaptive differences in responses to bacterial disease^44^. Taken together, this provides an avenue for questioning whether local populations are adapted to their native microbial environment.

In contrast to microbiomes, relatively few studies have characterized cnidarian viromes. To date, viromes have been described for a limited number of taxa, including several coral species^45–47^, *Hydra*^48^, *Nematostella* ^49^, and *Exaiptasia* ^50^. These studies, along with an additional analysis of 417 publicly available invertebrate RNA-Seq datasets that included more than 15 cnidarian species ^16^, examined viromes using previously published cnidarian sequencing data, although these samples were not collected specifically for virome analysis. Recent findings also indicate that viruses impact symbiotic dinoflagellates of corals^51^, suggesting a role for viruses in cnidarian symbiosis^52^. However, how viromes vary at the population level and how virus communities may differ between natural and laboratory conditions remains unexplored in cnidarians.

The sea anemone *N. vectensis*, a model cnidarian for both laboratory and field studies, is a prime system to examine environmental (i.e., virome and microbiome) and population-related questions. *N. vectensis* has a natural range along the Atlantic coast of North America, spanning from Nova Scotia (NS) to Florida (FL), with the widely used laboratory strain originally collected from Maryland (MD). A number of studies have revealed that populations (i.e., genotypes) along the North American Atlantic coast possess distinct microbiomes that are maintained over time under laboratory conditions^37^ and in natural habitats^53,54^. Additionally, studies have shown that many factors, such as the location of origin (i.e., genotype^37^), changes in diel light cycle^55^, life history stage^56,57^, and body region^58^, impact the bacterial composition of *N. vectensis*. Furthermore, comparisons between laboratory strains and natural populations along the North American Atlantic coast revealed differences in bacterial diversity between laboratory animals and wild populations, with substantial variation in associated bacteria among populations ^53,54^. Together, these results demonstrate that bacterial associations in *N. vectensis* are complex and vary across genotypes. In contrast, viral diversity in *N. vectensis* remains relatively understudied. A recent analysis of publicly available *N. vectensis* RNA-seq datasets identified a core virome and revealed variation in viromes across laboratory populations cultured in different laboratories^49^. Yet, how the *N. vectensis* virome differs across geographic populations and genotypes remains unexplored.

In this study, we used a mesocosm experiment to examine how individuals from six *N. vectensis* populations respond to natural microbial and viral communities from an estuary where this species occurs. By integrating microbiome, virome, and host gene expression data, we tested whether host genotype influences responses to a novel environment and whether these responses reflect population-level differences. We hypothesized that *N. vectensis* is locally adapted to the viromes and microbiomes of its native environment and would therefore be more sensitive to “foreign” communities.

## Materials and Methods

### Nematostella vectensis Animal Culture

Six female *N. vectensis* clonal lines (i.e., genotypes) were cultured in the laboratory prior to the mesocosm field study. Individuals used for the propagation of these lines were originally collected from Crescent Beach, Nova Scotia (NS), Saco River, Maine (ME), Wallis Sands, New Hampshire (NH), Sippewissett, Massachusetts (MA), Georgetown, South Carolina (SC), and near St. Augustine, Florida (FL) (see Supplemental Table 1 for GPS coordinates). *N. vectensis* clonal lines were maintained at the standard laboratory salinity of 15 ‰ in separate bowls for each population. Animal bowls were partially filled with filtered sediment (Onyx Sand; Seachem, product number 3505). Water changes and *Artemia salina* (brine shrimp) feedings were conducted every two days. Prior to the field study, all animals were shipped to the University of New Hampshire (UNH) and starved for two days. Animals were maintained at 15 ‰ in separate bowls until the start of the experiment.

### Mesocosm Field Study

Mesocosms were deployed at the University of New Hampshire (UNH) Jackson Estuarine Laboratory (Durham, New Hampshire, USA) in June 2022. In total, we conducted five replicate mesocosms (Figure 1A). Each mesocosm consisted of a plastic bin (58.5 cm L x 38.1 cm W x 33 cm H) containing a glass dish (9.5 in W x 16.38 in L x 3.25 in H) to attach animal tubes for each clonal line and to weigh them down (Supplemental Figure 1). Animal tubes were constructed from 50 mL conical tubes (Falcon, product number 352098) by removing the tips. Each end had a 100 µm strainer (Greiner, product number 542000), silicone-glued to the inside of the tube to contain animals and allow water flow. Each animal tube contained one *N. vectensis* genotype (i.e., clonal line) with six individuals. Animal tubes were attached to the glass dish using Velcro and bungee cords to keep them in place at the bottom of the mesocosm. Animal tube order was randomized for each mesocosm. Each mesocosm was filled with natural estuarine water collected from the estuarine lab using a pump connected to the inlet, and then bag-filtered to remove larger debris (200 µm; Pentair Industrial, product number KO200K2S). Additionally, each mesocosm contained a 30W, 550 GPH water pump (Freesea, product number FS-030R) to maintain gently flowing water throughout the mesocosm. Mesocosm water was replaced every two days with fresh bag-filtered estuarine water throughout the experiment. Whole animals were collected at two time points for sequencing: first, at the start of the experiment (T0), and then on day 14 (T14), marking the conclusion of the study.

**Figure 1.**
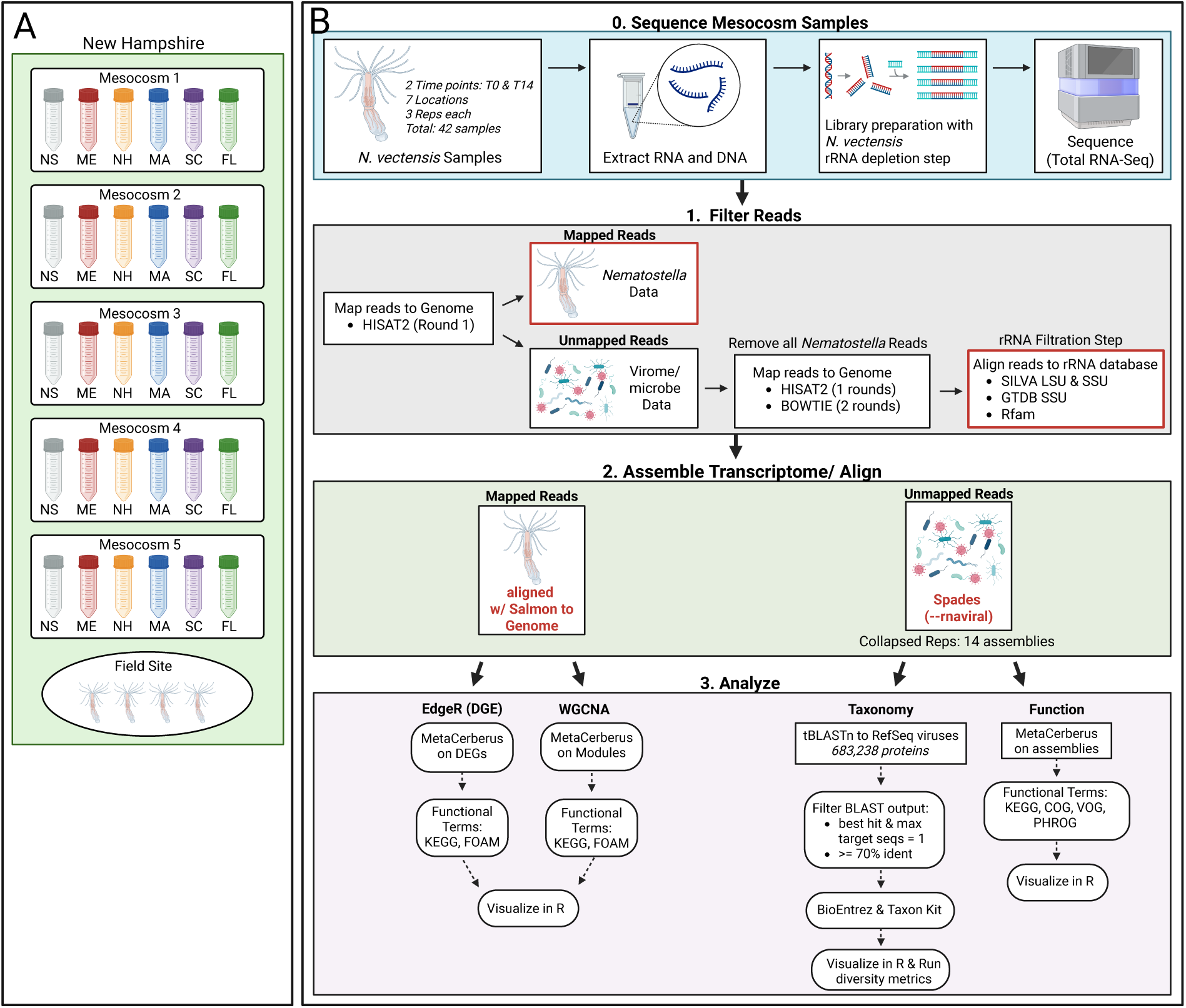
Mesocosm design and virome pipeline. (**A**) Cartoon depiction of mesocosm design, which included five replicate mesocosms, where each mesocosm contained six separated genotypes (Nova Scotia, NS; Maine, ME; New Hampshire, NH; Massachusetts, MA; South Carolina, SC; Florida, FL). Animals were collected at two time points: T0, prior to entering the mesocosms, and T14, 14 days after mesocosm exposure. A nearby natural population was sampled on the same collection days. This study was conducted in New Hampshire. See Supplemental Figure 1 for the actual mesocosm setup. (**B**) General virome pipeline from extraction through data analysis visualizations. After sequencing, samples underwent a read filtration step where we separated Host reads (*N. vectensis*) and presumably viral reads. We then assembled viral transcriptomes and performed taxonomy and functional term analyses. The host reads were aligned to the *N. vectensis* genome, and a differential gene expression analysis and WGCNA were performed.

In addition to the experimental mesocosm samples, samples from the nearest natural population of *N. vectensis* in New Hampshire were collected to compare anemones from natural field environments with lab-cultured animals at the same time points (T0 and T14). The natural population (referred to as field) was sampled from a marsh across from Wallis Sands Beach (Rye, New Hampshire, USA; 43.028656, –70.730775). Collections were made by scooping marsh pond mud onto a window screen and placing wild *N. vectensis* into a 50 mL tube filled with estuarine water for transport to the Jackson Estuarine Laboratory. All sampled animals (i.e., mesocosm and field) were flash frozen and stored at -80 °C. Samples were then shipped to the UNC Charlotte laboratory on dry ice and immediately stored at -80 °C until extraction.

### Extraction of Nucleic Acids and Total RNA Sequencing

Both RNA and DNA were extracted from the same individual anemones using the Qiagen All-Prep kit (product number 80204) with minor alterations to the manufacturer’s protocol. In step 2, 600 µl of RLT plus buffer was added (instead of the recommended 350 µl), and the whole animal was homogenized using a 3 ml syringe (BD Plastipak, product number 309577) to improve extraction efficiency. Samples were placed on ice for 10 minutes and then centrifuged as recommended in step 4. For total RNA purification, 350 µl of 70 % ethanol was added (instead of the recommended 600 µl), and the remainder of the protocol was completed according to the manufacturer’s specifications. Samples were quantified using Qubit High Sensitivity RNA reagent (Thermo Fisher Scientific, product number Q32855), NanoDrop (Thermo Fisher Scientific), and TapeStation High Sensitivity RNA assay (Agilent, product number 5067-5579). All samples had RIN values above 7.0. Libraries were created using 500 ng of total RNA using the Tecan Universal Plus Total RNA-Seq library kit with an *N. vectensis* rRNA depletion step (Tecan, product numbers 0361-24, 0370-24, S02713). Libraries were sequenced on the NovaSeq X Plus platform (125-bp PE, Illumina, performed by Admera Health, Plainfield, New Jersey, USA).

### *N. vectensis* Virome Read Processing and Analysis

A custom pipeline was developed to identify sequence reads that primarily originated from viruses (Figure 1B). First, Trimmomatic ^59^ and FastQC ^60^ were run on raw reads to remove adaptors and assess read quality. Using custom Python scripts (available at GitHub https://github.com/sydney-birch/Nematostella_Mesocosm_virome), these trimmed reads were aligned to the *N. vectensis* genome ^61^ with HISAT2 ^62^ to filter out host reads from the anemone. SAMtools ^63^ was then used to generate fastq files for reads that did not map to the reference genome (GCA_932526225.1; Fletcher and Pereira 2023). To ensure anemone reads were removed, an additional round of mapping with HISAT2 and a round of mapping with Bowtie2 using high sensitivity (--very-sensitive-local) ^64^ were performed until no reads mapped to the *N. vectensis* genome. Next, an rRNA filtration step was implemented where all unmapped reads were aligned to an rRNA database comprised of SILVA LSU and SSU, GTDB SSU, and Rfam sequences ^65–67^ to remove any eukaryotic and prokaryotic reads, leaving presumably viral reads with potentially unmapped or unidentified microbial reads. An additional filtration step was run against the RefSeq NR database ^68^, but no reads mapped to this database; therefore, the rRNA filtration dataset was used moving forward and will be referred to as “presumably viral reads”. Next, the presumably viral reads were assembled at each location and time point, creating a total of 14 assemblies (i.e., three replicates assembled for each location: NS, ME, NH, Field, MA, SC, FL at T0 and T14), using Spades rnaviral ^69^.

The 14 presumably viral assemblies were used to investigate viral taxonomy and function. To investigate taxonomy, a tBLASTn ^70,71^ was performed, querying the RefSeq virus protein database, comprised of 683,238 viral proteins ^72^, against the 14 presumably viral assemblies. The tBLASTn run included the best-hit flag, max target seqs equal to 1, and an e-value of 1e-5. The BLAST output was filtered to include only proteins (hits) with 70 % identity or greater. This decision was based on comparisons of lower percentage thresholds. The 70 % identity threshold produced a T0/T14 read ratio like that of the total BLAST hits and had a manageable number of protein hits to analyze (i.e., an average of 44,244 hits compared to an average of 786 hits). Next, a custom Python script using BioEntrez ^73^ was used to retrieve Taxon IDs from the BLAST output accession IDs. The Taxon IDs were then used with NCBI TaxonKit ^74^ to retrieve the full taxonomic information (i.e., kingdom to strain) for all proteins in the dataset.

The viral genetic composition was obtained from the International Committee on Taxonomy of Viruses (ICTV) Virus Metadata Resource (VMR) file ^75^. Visualizations of counts and diversity metrics were completed in RStudio using the vegan package. MetaCerberus, an automated pipeline that processes sequencing data through various functional databases ^76^, was used to investigate viral function. The 14 viral assemblies were input to the program with the Prodigal flag selected to find protein-coding open reading frames, and all databases (i.e., VOG, PHROG, KEGG, FOAM, COG, and CAZy) were selected. Visualizations were completed in RStudio.

Lastly, this dataset was compared with the 10 core qPCR-verified viruses previously identified in *N. vectensis* ^49^. Here, a BLAST search of the qPCR-verified viruses was completed against the 14 presumably viral assemblies using the following parameters: max target seqs set to 50 and the e-value set to 1e-5. Visualizations of counts were completed in RStudio. All bioinformatic processing and analyses are available on GitHub: https://github.com/sydney-birch/Nematostella_Mesocosm_virome.

### 16S rRNA Gene Sequencing and Analysis

The hypervariable regions V1-V2 of bacterial 16S rRNA genes were amplified for each sample (i.e., five replicates for each location and time point) using the following forward (5’-CCTACGGGNGGCWGCAG-3’) and reverse primers (5’-GACTACHVGGGTATCTAATCC-3’) ^53^ with the DNA extracted from the Qiagen All-Prep kit (described above). For the first PCR, 100 ng of template DNA was added to a 25 μL reaction and run on an Eppendorf thermal cycler (Eppendorf, Mastercycler Nexus). Each reaction contained 12.5 µl of Kapa HiFi HotStart Ready Mix (2x) (Roche, product number 09420398001), 0.75 µl of forward primer, 0.75 µl of reverse primer, 5 µl of template DNA, and 6 µl of DNase-free water. The PCR was conducted with the following cycling conditions (98 °C – 30s, 30 x [98 °C – 9s, 55 °C – 60s, 72 °C – 90s], 72 °C – 10 min). Nextera DNA index primers and Illumina sequencing adapters (Set C; FC-131-2003) were attached to the primers in the second PCR. The second PCR consisted of a 50 µL reaction containing 25 µL of Kapa Ready Mix (2x), 5 µL of Nextera XT index i7 primer, 5 µL of Nextera XT index i5 primer, 10 µL of DNase-free water, and 5 µL of 16S rRNA sample. The second PCR was conducted with the following cycling conditions (95 °C – 3 min, 8 x [95 °C – 30s, 55 °C – 30s, 72 °C – 30s], 72 °C – 5 min). Samples were then subjected to bead cleanup and quantified using TapeStation (DNA High sensitivity D1000; Agilent, 5067-5584 and 5067-5585) and Qubit (High sensitivity dsDNA; Thermo Fisher Scientific, Q33230). Paired-end samples (2x250) were sequenced on the Illumina NextSeq 2000 platform (Admera Health, Plainfield, NJ, USA).

QIIME2 v2024.2 ^77^ was used to process and analyze 16S rRNA gene sequences, with five replicates per sample. Demultiplexed paired-end reads were imported with the Casava 1.8 paired-end demultiplexed plug-in ^77^. Sequence quality filtering, denoising, chimera removal, and feature table construction were implemented using the DADA2 plugin ^78^. Next, a phylogenetic tree of the amplicon sequence variants (ASVs) was generated for use in diversity analyses using MAFFT ^79^ and FastTree2 ^80^. A sampling depth of 61,745 reads was selected to run the core metrics for the phylogenetic analysis. Two replicates from Field T14 with low read counts were excluded, leaving three replicates for this sample’s timepoint to retain more microbial features across the dataset. Alpha diversity metrics (Faith phylogenetic diversity, evenness metrics, Shannon index) and beta diversity metrics (Bray–Curtis, Jaccard, Weighted-Unifrac, Unweighted-Unifrac) were estimated using the q2-diversity plug-in, which utilized the rooted phylogenetic tree generated from the core metrics. Alpha diversity statistical tests employed Kruskal-Wallis ^81^, examining all groups and then performing pairwise comparisons for the following factors: time point (T0, T14), genotype (NS, ME, NH, Field, MA, SC, FL), Northern genotypes (NS, ME, NH) vs Southern genotypes (MA, SC, FL), and the time points and genotypes combined (NS-T0, NS-T14, NH-T0, etc.). Principal Coordinate Analyses (PCoAs) were estimated for the beta-diversity metrics using emperor from the q2-diversity plug-in. The diversity metrics were then exported to RStudio for visualization. The taxonomic analysis was completed using a Greengenes classifier (2022.10.backbone.full-length.nb.qza) to assign taxonomy to ASVs ^82^, and the results were visualized in RStudio.

### *N. vectensis* RNA read processing and analysis

Host sequence reads from the anemone were identified by aligning sequence reads to a *N. vectensis* reference genome (see virome section above, Figure 1B). Host reads were quantified using Salmon ^83^. Two analyses were performed using the Salmon-quantified reads.

First, a differential gene expression analysis was conducted using EdgeR ^84^ to perform pairwise comparisons between time points for each genotype. Transcripts were required to have at least 10 counts-per-million to be included in the analysis ^84^. Library size normalization was completed using the calcNormFactors function, which employs a trimmed mean of M-values (TMM) method. Dispersion estimation was conducted with a quantile-adjusted conditional maximum likelihood (qCML) method. Differentially expressed genes (DEGs) were identified at a p-value of 0.01 using the Benjamin-Hochberg correction. The DEGs were imported to MetaCerberus ^76^ to obtain functional annotations using the eukaryotic module (i.e., KEGG and FOAM databases). Functional groups and heatmaps of differentially expressed genes were visualized in RStudio. Additionally, expression of 56 immune-related *N. vectensis* genes ^85^ was examined by generating a heatmap of normalized expression in RStudio.

Second, a weighted gene co-expression network analysis (WGCNA) ^86^ was performed using the Salmon-quantified reads. A DESeq2 ^87^ dataset was created without specifying a model, since the purpose was not to perform differential expression analysis but to normalize and transform our data. Normalization and transformation were completed with the ‘vst()’ function from the DESeq2 package. The ‘pickSoftThreshold()’ function from the ‘WGCNA’ package was used to identify the WGCNA power parameter, which affects the number of modules identified. A power threshold above 0.80 was selected, as recommended ^88^. The WGCNA was run, and two model matrices were designed: one to examine the effect of time point (T0 vs T14) and one to examine the effect of genotype region (North vs South vs Field). The geographic locations of genotypes were separated based on Cape Cod, MA, USA. For each model matrix, a linear model was run on each module using Limma, and multiple-testing corrections were then applied. The resulting statistics were used to generate figures for both models (time point and genotype region). Module genes were then analyzed using MetaCerberus to identify KEGG and FOAM functional groups, which were visualized in RStudio. All bioinformatic processing and analyses are available on GitHub: https://github.com/sydney-birch/Nematostella_Mesocosm_virome.

## Results

### Virus Taxonomy and Diversity

To determine which viral groups *N. vectensis* genotypes associate with before and after mesocosm exposures, relative to the natural field population, we first examined the viral genomic composition (Figure 2A). Across all conditions, dsDNA viruses predominated. Following 14 days of mesocosm exposure (T14), the relative abundance of dsDNA viruses increased significantly across genotypes (p = 0.0009). We also detected a significant increase in ssRNA(-) viruses (p=0.024) and unknowns (p=0.003), while no dsRNA viruses were recovered (Figure 2A).

**Figure 2.**
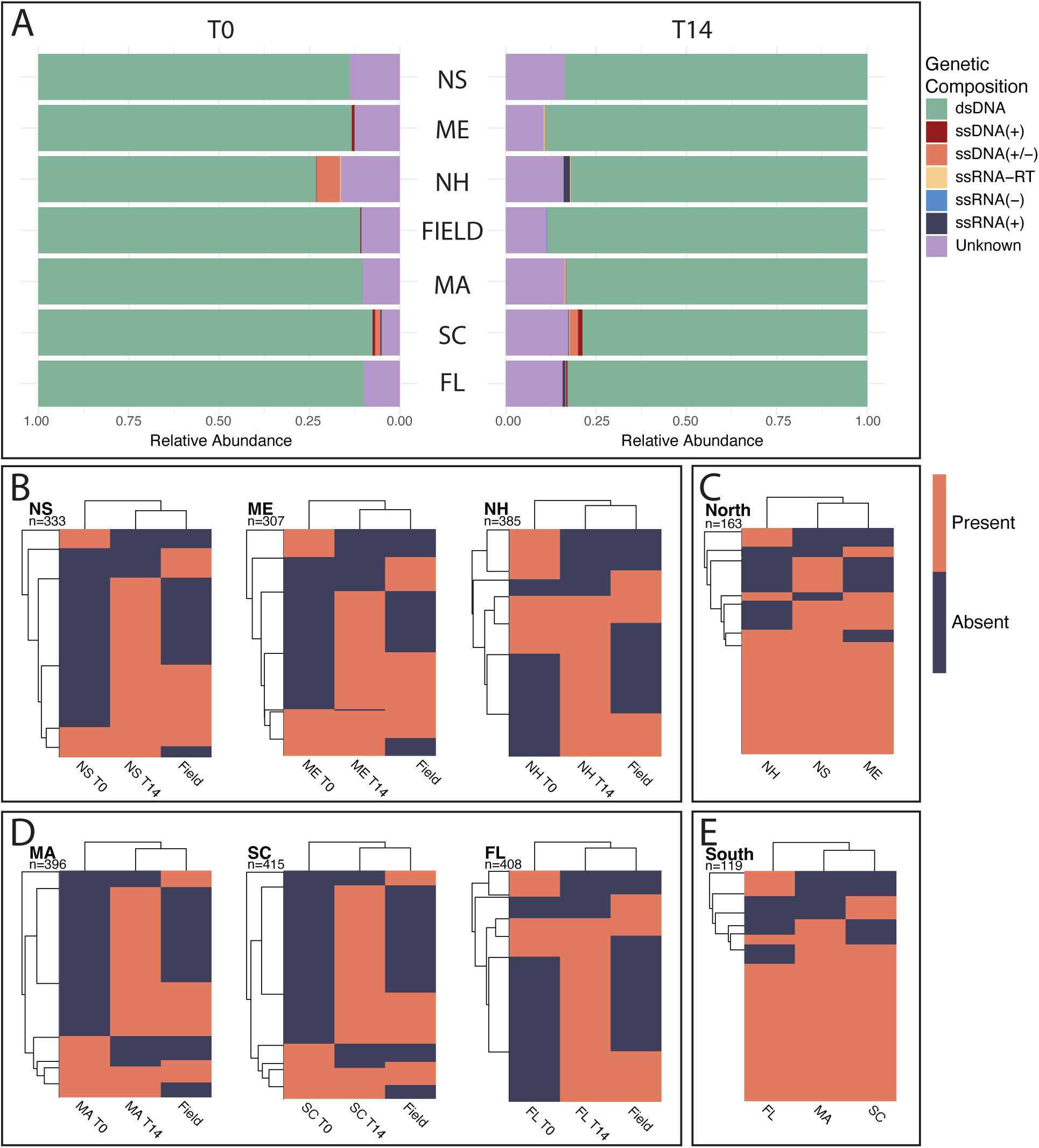
Types of viruses and viral genera associated with *N. vectensis* genotypes. (**A**) Mirrored bar plot of the relative abundance of the genetic composition of viral taxa found at T0 and T14 across genotype locations. Viral genetic compositions were retrieved from the International Committee on Taxonomy of Viruses (ICTV) Virus Metadata Resource (VMR). (**B-E**) Heatmaps of the presence and absence of viral genera in (**B**) northern genotypes and (**D**) southern genotypes. Each heatmap presents a comparison of a specific genotype at T0 and T14 with the natural population (Field). Each row represents a single viral genus; the total number of genera in the heatmap is located below the genotype abbreviation. Presence of a viral genus is indicated in orange, and absence is indicated in dark blue. (**C**) The union between each northern genotype (i.e., NS, ME, NH) at T14 and the natural population. This comparison examines the shared T14 (newly accumulated) viruses of northern genotypes with the natural population. (**E**) The union between each southern genotype (i.e., MA, SC, FL) at T14 and the natural population. This comparison examines the shared T14 (newly accumulated) viruses of southern genotypes with the natural population. See Supplemental Figure 2 for viral diversity metrics.

Next, we compared viral taxa between genotypes and field-collected anemones across time points by generating presence/absence heatmaps to visualize overlap in viral genera (Figure 2B-E). Both northern (i.e., NS, ME, NH; Figure 2B) and southern (i.e., MA, SC, FL; Figure 2D) genotypes showed a lack of viral taxa prior to entering the mesocosm at time point 0 (T0) compared to the natural field population and the T14 animals. Additionally, across all genotypes, we observed an increase in the number of viral taxa (i.e., viral load) at T14 following mesocosm exposure, with southern genotypes showing a higher overall viral load (p=0.0105). When comparing the T14 animals to the natural field population, we observed genotype-specific gains in viral taxa that were absent from the field population or the T0 animals.

Furthermore, alpha-diversity metrics revealed that species richness (*p*=0.00155), Shannon diversity (*p*=0.00048), and species composition (*p*=0.003) were significantly increased at T14, with southern genotypes having greater viral diversity (Supplemental Figure 2). Beta-diversity was assessed using an NMDS (Supplemental Figure 2), which showed substantial variation among T0 genotype samples and tight clustering of T14 genotype samples with the natural population. However, both T0 and T14 genotype samples are regionally separated (north vs. south), with T14 genotypes distinct from the natural population (Supplemental Figure 2).

Next, we examined the union (shared viruses) between the natural population and the T14 animals to determine whether the northern (Figure 2C) and southern (Figure 2E) genotypes in the mesocosms shared the same viral taxa as the natural field population. Both northern and southern populations predominantly associated with the same viral taxa across the respective genotypes, with southern genotypes sharing more viral taxa across all three genotypes (74 viruses shared across FL, SC, and MA) than northern genotypes (54 viruses shared across NS, ME, and NH). Furthermore, we found that 13 taxa were shared across all locations and time points, including Alphabaculovirus, Avipoxvirus, Betaentomopoxvirus, Tupanvirus, Betabaculovirus, Chlorovirus, Rheavirus, Marseillevirus, Prasinovirus, Oceanusvirus, and unclassified viruses in Phycodnaviridae, Caudoviricetes, and Marseilleviridae. Additionally, the two most prevalent viruses in the dataset belonged to the genera Alphabaculovirus, a large DNA virus that infects invertebrates ^89^, and Tequatrovirus, a bacteriophage that infects Gram-negative bacteria using a lytic cycle to reproduce ^90^.

Lastly, we compared our viral dataset with the 10 qPCR-verified core viruses from a previous *N. vectensis* virome survey ^49^ (Supplemental Figure 3). The natural population (field) did not possess any of the 10 verified viruses, which included three *Artemia* viruses and seven other invertebrate-associated viruses. Additionally, all time points for each genotype, except the field population, harbored the three *Artemia* viruses (Shahe yuevirus-like virus 1, Wuchan romanomermis nematode virus 2, and Beihai sesarmid crab virus 3), but only the Maine genotype harbored all seven other viruses at both time points. The other five anemone genotypes at both time points contained only two of the seven viruses: Hubei Sobemo-like virus 41 and Sanxia water strider virus 10.

### Viral Functional Terms

To examine the annotated viral proteins and their potential functions across each *N. vectensis* genotype (i.e., clonal line) and time point, we assembled viral transcriptomes and analyzed them using MetaCerberus ^76^ (Figure 3). We first assessed the number of viral proteins identified by MetaCerberus and found an increase in viral protein count at T14 compared with the initial time point (T0), across all genotypes (*p* = 0.00087; Figure 3A). The natural population showed similar counts at T0 and T14, as expected (Figure 3). Additionally, the southern genotypes (i.e., MA, SC, FL) had higher viral protein counts at T14 than the northern genotypes (i.e., NS, ME, NH; *p*=0.002; Figure 3A). This same pattern occurred for the number of virus orthologous group (VOG) ID counts (Figure 3B; *p*=0.00028) and prokaryotic virus remote homologous group (PHROG) ID counts (Figure 3C; *p*=0.001).

**Figure 3.**
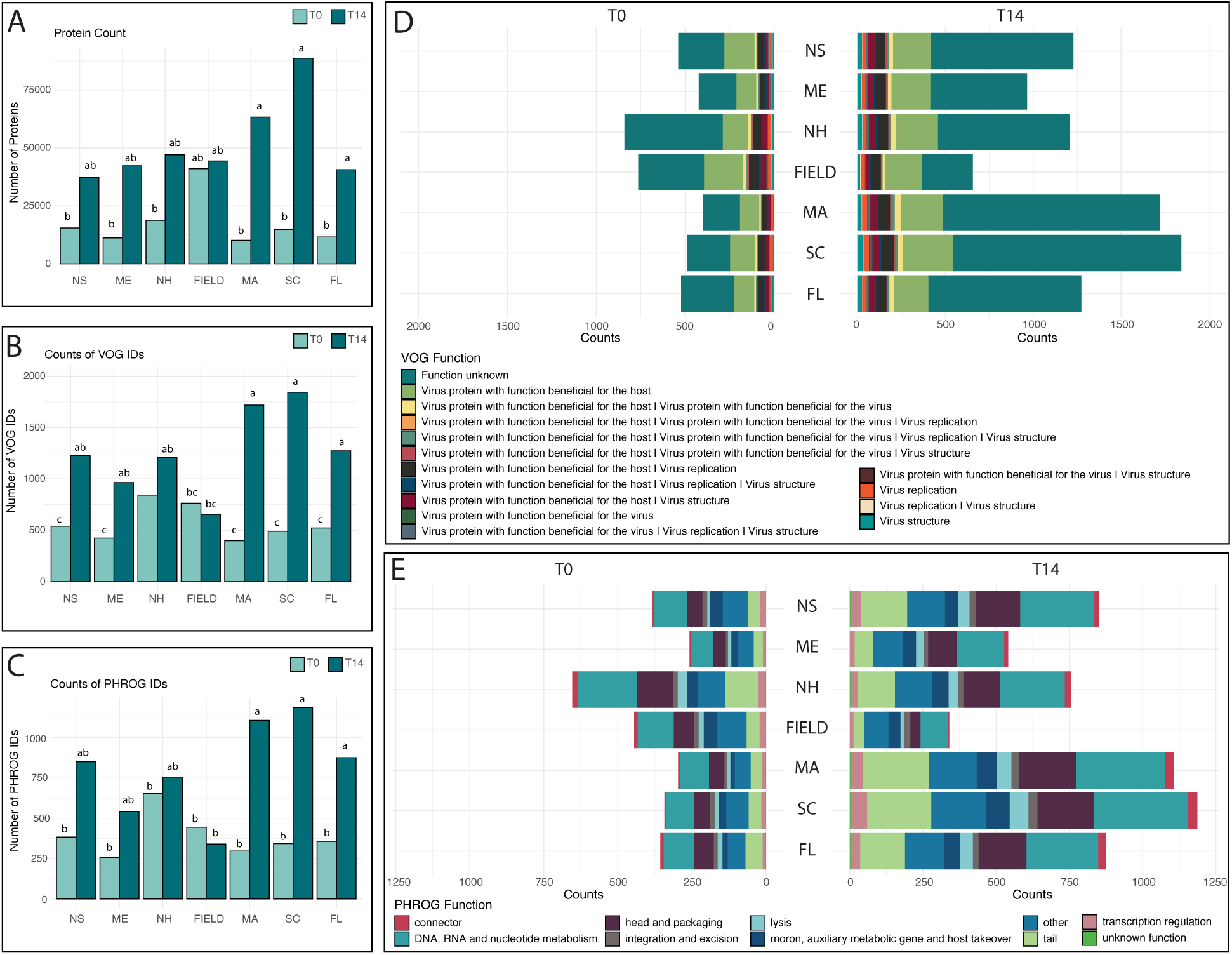
Viral functional terms identified at each time point for each *N. vectensis* genotype. (**A**) The number of viral proteins identified from each *N. vectensis* genotype and time point using MetaCerberus. Light teal represents T0 and dark teal represents T14. Significance groups are derived from a Tukey comparison of timepoints of genotype-region (i.e., North T0, North T14, etc.). (**B**) The number of Virus Orthologous Group (VOG) term IDs identified for each *N. vectensis* genotype and time point. (**C**) The number of prokaryotic virus remote homologous groups (PHROG) term IDs identified for each *N. vectensis* genotype and time point. (**D**) Mirrored bar plot of VOG functional term counts across genotypes and timepoints. (**E**) Mirrored bar plot of PHROG functional term counts across genotypes and timepoints. See Supplemental Figure 4 for functional term statistics from VOG and PHROG databases. Additionally, see Supplemental Figure 5 for KEGG mirrored bar plot of functional terms.

Subsequently, we examined the functional terms associated with the annotated viral proteins, focusing predominantly on the virus (VOG) and phage (PHROG) databases. The following VOG functions were significantly increased in ID counts at T14: function unknown (*p*=0.0004), virus proteins with functions beneficial for the host (*p*=0.0013), and viral structure (*p*=0.002) and replication (*p*=0.015) proteins (Figure 3D; Supplemental Figure 4). The following PHROG functional terms also displayed a significant increase in ID counts at T14: lysis (*p*=0.0007), tail (*p*=0.003), head and packaging (*p*=0.0009), and connector (*p*=0.001; Figure 3E; Supplemental Figure 4). Furthermore, we investigated these VOG and PHROG terms to determine how term counts varied across regions and time points (Supplemental Figure 4). For both term databases, these terms all showed considerably more functional terms present in the southern genotypes at T14 (i.e., MA, SC, FL). Finally, KEGG terms had a significant increase in ID counts at T14 for genetic information processing (p=0.0002), infectious disease: viral (p=6.1e-05) and bacterial (p=6.82e-05), immune system (p=0.00023), and environmental adaptation (p=2.62e-05; Supplemental Figures 5 and 6).

### Bacterial Diversity

To investigate the bacterial associations of *N. vectensis* genotypes across locations and time points, we performed a 16S rRNA analysis using QIIME2 ^77^. We first examined the alpha- and beta-diversity of all ASVs. The alpha-diversity revealed a significant increase in Faith PD (*p*=0.015; Figure 4C), Shannon Diversity (*p*=0.044; Figure 4B), and evenness (*p*=0.021; Supplemental Figure 7C) in southern genotypes compared to northern genotypes, with genotype location being a significant factor (Faith PD *p*=0.027; Evenness *p*=0.019, Shannon Diversity *p*=ns). Additionally, only the FL and NH genotypes showed a significant increase in bacterial diversity after mesocosm exposure (Shannon Diversity; Figure 4B; *p*=0.008; Faith PD; Figure 4C; *p*=0.014). Beta diversity was assessed with Bray-Curtis (Figure 4D), Jaccard (Supplemental Figure 7B), and unweighted UniFrac (Supplemental Figure 7A). All three PCAs reveal clustering of samples by time point. Additionally, SC and FL clustered together, with MA clustering closer to the northern genotypes (Figure 4D), in contrast to the virome, where MA clustered with southern genotypes.

**Figure 4.**
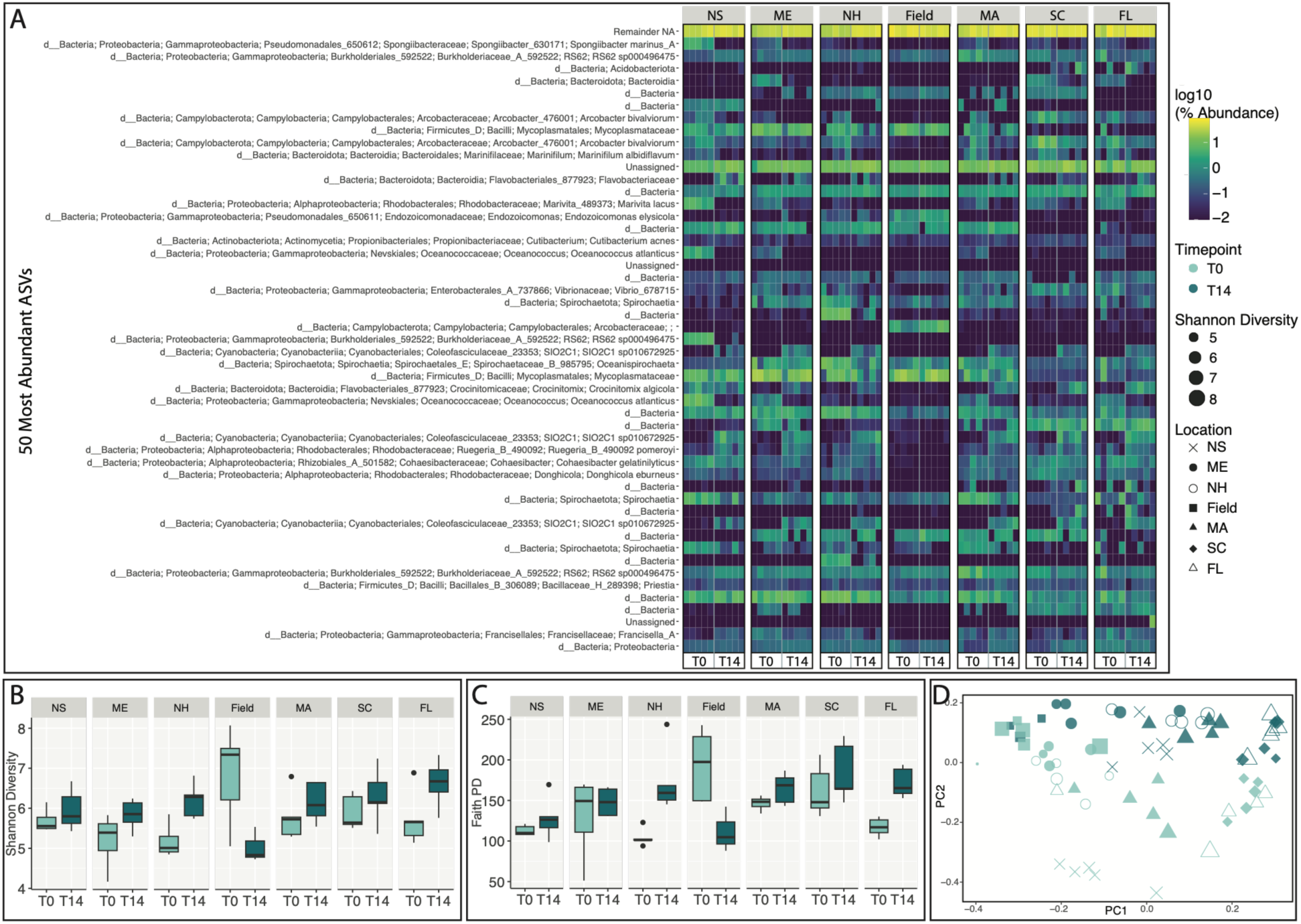
Most abundant bacterial features and bacterial diversity statistics. (**A**) Heatmap of percent abundance of the top 50 most abundant ASVs associated with each *N. vectensis* genotype and timepoint. Yellow represents high abundance, and dark blue represents low abundance. Boxplots of alpha diversity metrics (**B**) Shannon diversity and (**C**) Faith PD across each genotype location and time point. (**D**) Bray-Curtis PCA assessment of beta diversity, where the size of the symbol correlates to Shannon diversity. Keys for time point colors and location symbols are to the right. See Supplemental Figure 6 for additional alpha and beta diversity plots.

Next, we examined the 50 most abundant ASVs (Figure 4A) and found that genotypes associated with bacterial ASVs in diverse patterns. Generally, the lab-reared anemones did not become more similar to the field population regardless of genotype. We found examples of different abundance patterns for the top 50 most abundant taxa, such as those with approximately the same abundance across all time points and locations (Burkholderiales_592522, *Cutibacterium acnes*, *Oceanispirochaeta*, and *Priestia*), increased abundance of particular bacterial ASVs at T14, excluding the field population (*Flavobacteriaceae*, Cyanobacteria spp., *Crocinitomix algicola*, and *Rhodobacteraceae* spp.), and the inverse, with a higher proportion at T0, excluding the field population (*Marivita lacus* and *Oceanococcus atlanticus*). Additionally, we identified cases in which all time points and genotype locations had higher levels of a particular bacterium than the field population (*Arcobacter bivalviorum*, Vibrio_678715, *Cohaesibacter gelatinilyticus*, and *Donghicola eburneus*), whereas only one instance showed the field population higher than the lab-cultured anemones (Arcobacteraceae).

### Bacterial Taxonomic Overlap

Next, we examined the relative abundance of all bacterial phyla identified across genotype locations and time points (Figure 5A). We identified an increase in cyanobacteria and Bacteroidota at T14 across all genotype locations, except for field populations. In contrast, members of the phylum Spirochaetota decreased at T14 across genotype locations except in field populations. Additionally, we found longitudinal differences in the bacterial phyla present. Firmicutes were more abundant in northern populations, including MA (i.e., NS, ME, NH, MA), and Desulfobacterota were more abundant in field and southern T0 populations (i.e., MA, SC, FL). Lastly, Proteobacteria showed less consistent patterns, with abundance varying by genotype, location, and time point.

**Figure 5.**
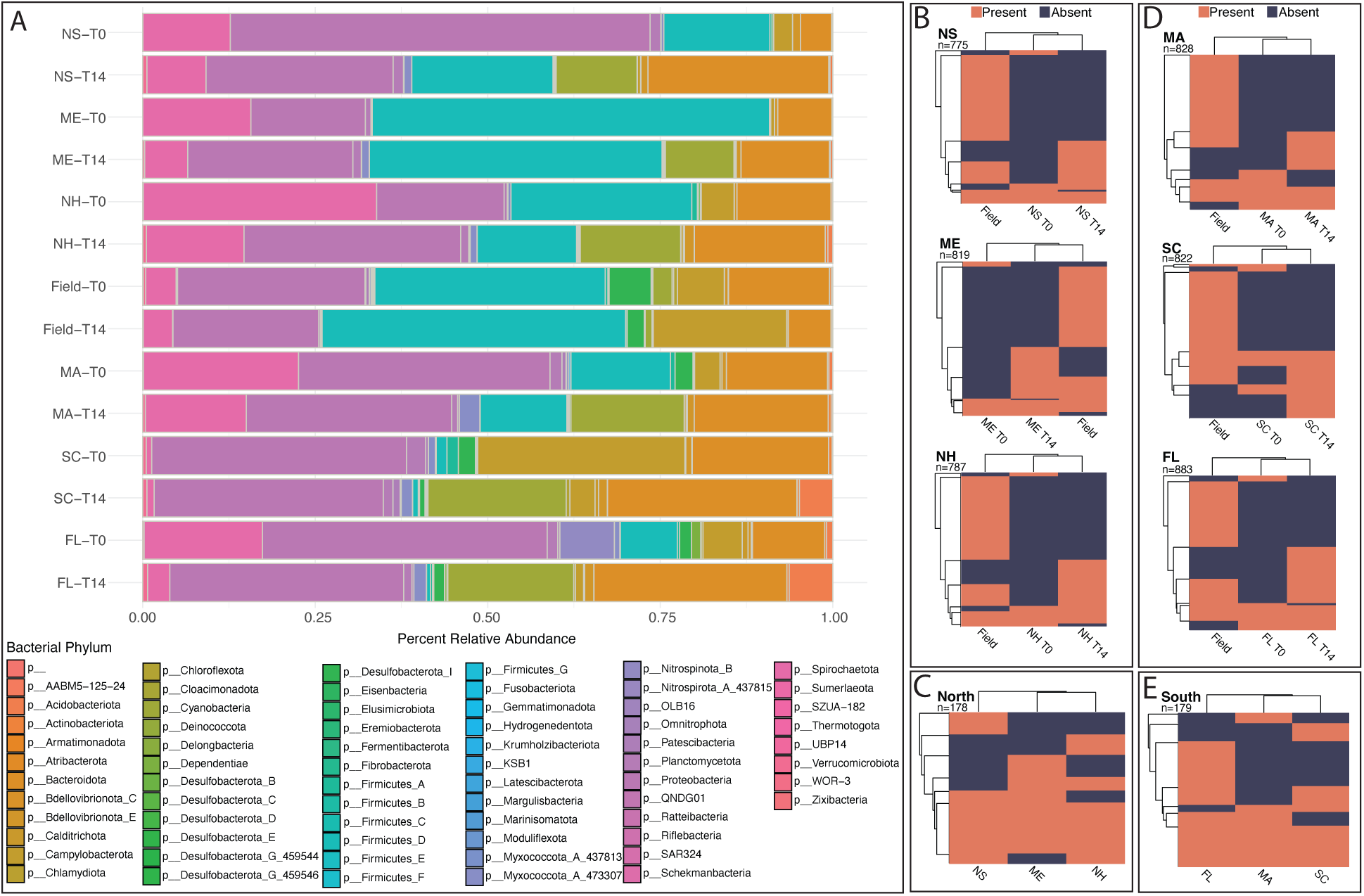
Bacterial phyla and bacterial genera associated with N. vectensis genotypes. (**A**) Percent relative abundance of all bacterial phyla found in each *N. vectensis* genotype and time point. (**B-E**) Heatmaps of the presence and absence of bacterial genera in (**B**) northern genotypes and (**D**) southern genotypes. Each heatmap presents a comparison of a specific genotype at T0 and T14 with the natural population (Field). Each row represents a single bacterial genus; the total number of genera in the heatmap is located below the genotype abbreviation. Presence of a bacterial genus is indicated in orange, and absence is indicated in dark blue. (**C**) The union between each northern genotype (i.e., NS, ME, NH) at T14 and the natural population. This comparison examines the shared T14 (newly accumulated) bacteria of northern genotypes with the natural population. (**E**) The union between each southern genotype (i.e., MA, SC, FL) at T14 and the natural population. This comparison examines the shared T14 (newly accumulated) bacteria of southern genotypes with the natural population. See Supplemental Figure 2 for viral diversity metrics.

Subsequently, we investigated how microbial genera associated with *N. vectensis* genotypes between time points and the natural field population (Figure 5 B-E). Here, we aimed to determine whether the microbial genera associated with anemones of different genotypes were the same across time points and whether these genera differed from those in the natural population. Similar to the viral analysis, the northern (NS, ME, NH; Figure 5B) and southern (MA, SC, FL; Figure 5D) genotypes all exhibited an increase in the presence of bacterial genera from T0 to T14; however, inconsistent with the viral associations, more bacterial associations were identified in the field population rather than with the T14 animals. Genotype-specific gains between T0 and T14 were observed, as in the virome analysis; however, the percentage of bacterial associations gained was lower than that of viruses (i.e., an average of 16% gain in bacteria at T14 compared to an average of 36% gain in viruses at T14). When we examined the shared bacterial taxa between T14 genotypes and the natural population, northern genotypes (NS, ME, NH; Figure 5C) and southern genotypes (MA, SC, FL; Figure 5E) associated primarily with the same bacterial genera. Northern genotypes shared 60 bacterial genera, including genotype-specific shared genera, while southern genotypes shared fewer total bacterial genera (48), with FL having the most genotype-specific shared genera with the field population.

### Host Transcriptomics

To examine the transcriptional response by *N. vectensis* to the mesocosm exposure, we first performed a differential gene expression analysis (Figure 6A; Supplemental Figure 8). We examined the differentially expressed genes (DEGs) within genotypes between time points and found that the most differentially expressed transcripts were present in NS (469 DEGs) and ME (377 DEGs), while all other locations had a low number of DEGs (43-12 DEGs), with FL having the lowest (12 DEGs). Additionally, the NS and ME genotypes had the greatest number of immune- and antimicrobial-related DEGs at T14. We then analyzed the DEGs using MetaCerberus to obtain functional information from the KEGG database (Figure 6A). We observed an average of 11 terms associated with the immune system and infectious disease: bacterial and viral, with the highest counts in the NS and ME genotypes. However, the stronger responses were in signaling and cellular processes, genetic information processing, and signal transduction, which included terms such as cytokine receptors, pattern recognition receptors, and antimicrobial resistance genes. To further examine the potential expression of the host immune-related response, we interrogated our data with a previously curated *N. vectensis* gene set of immune-related genes ^85^, Supplemental Figure 9. There was a slight increase in gene expression in northern T14 genotypes in the upper clades of the heatmap, with the lowest expression pattern occurring in FL at T14 (Supplemental Figure 9). The two upper clades included genes involved in innate immunity in *N. vectensis*, including *cGAS*, *RLRa*, and *RLRb*, among others. Additionally, we found that among the *N. vectensis* immune-related gene set, only one gene (*GBP3-like*) was significantly differentially expressed in NS and NH.

**Figure 6.**
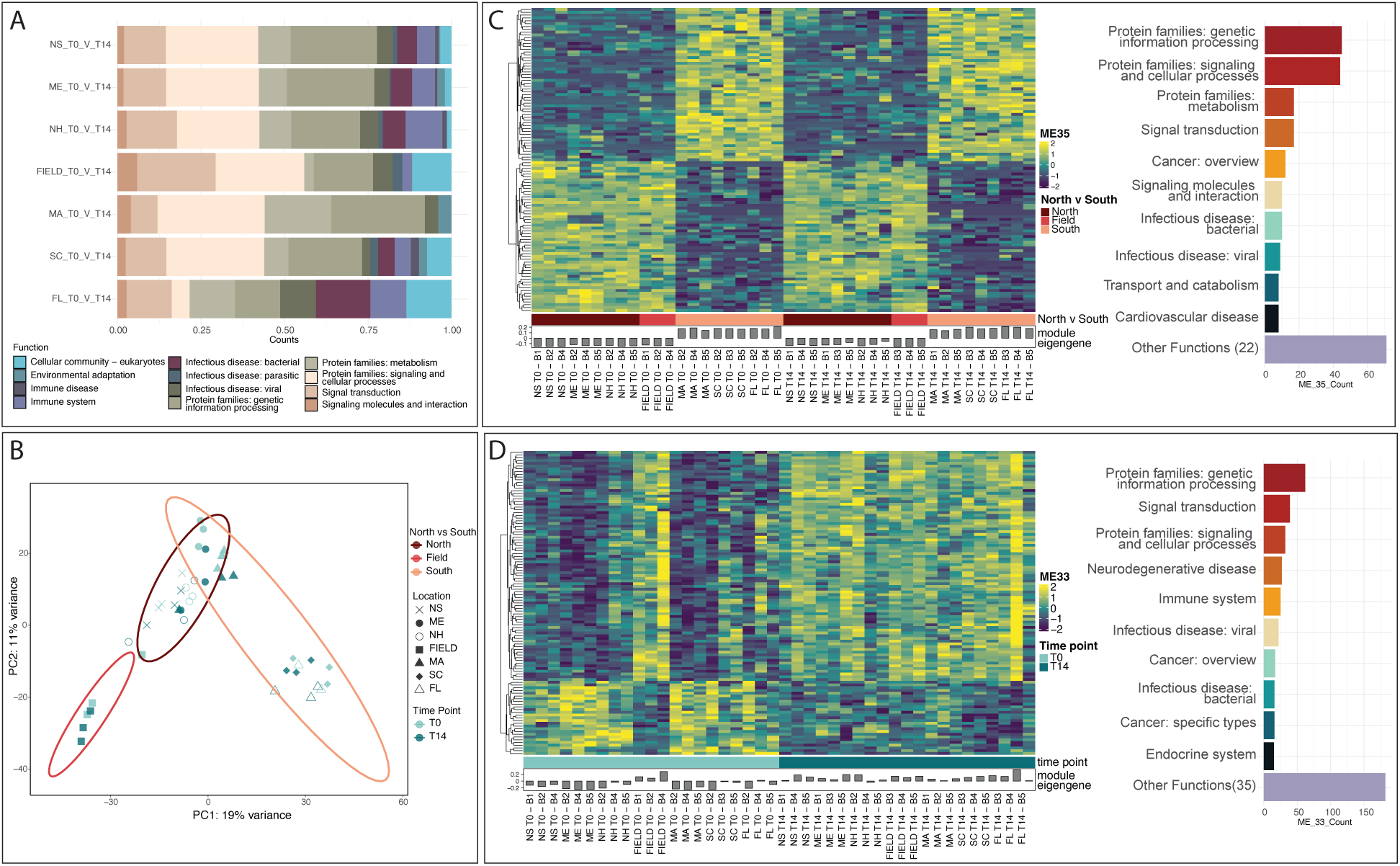
Differential gene expression and WGCNA of N. vectensis genotypes and timepoints. **(A)** Differential gene expression analysis. Differentially expressed genes (DEGs) were identified using the EdgeR package by comparing expression data between time points for each genotype and were then annotated into functional groups using the KEGG database. The stacked bar plot depicts the relative abundance of DEG counts within KEGG functional terms. See Supplemental Figure 7 for heatmaps of DEGs between time points for each genotype. (**B-D**) WGCNA analysis. **(B)** PCA of *N. vectensis* (host) expression data. Time point is indicated by color, the symbols represent genotype locations, and the ellipses identify clustering of genotype region (i.e., North [NS, ME, NH], South [MA, SC, FL], and Field). (**C**) The most significant eigengene module (Module 35) for the genotype-region comparison (i.e., North [NS, ME, NH], South [MA, SC, FL], and Field). The genes within the module were further annotated using the KEGG database. (**D**) The most significant eigengene module for the timepoint comparison (Module 33). The genes within the module were further annotated using the KEGG database.

Next, we conducted a PCA and WGCNA to identify correlations across all *N. vectensis* samples (Figure 6B-D). The PCA revealed that the natural population clustered separately from the experimental (mesocosm) samples. Among the mesocosm samples, the northern genotypes (NS, ME, NH) clustered together and were distinct from southern genotypes (MA, SC, FL), except for MA, which clustered with the northern genotypes (Figure 6B). There was a modest separation between time points.

WGCNA explored correlated gene expression for two design matrices: genotype-region (north vs. south) (Figure 6C) and time point (Figure 6D). The genotype-region model identified 34 significant eigengene modules (adjusted P-values) by examining the eigengene modules that showed the greatest differences between northern and southern genotypes. The most significant module from the genotype-region investigation was module 35 (ME 35; *p*=2.85e-22; Figure 6C). This module contained 107 genes and showed an inverse expression pattern between northern (NS, ME, NH) and southern (MA, SC, FL) genotypes, with the field population displaying a pattern similar to that of the northern genotypes. Among the genes in this module were homologs of the mammalian E3 ubiquitin proteins (bacterial recognition proteins; Cheng et al. 2016), integrin beta-6 (potential receptor for viruses; Heikkilä et al. 2009; Gianni et al. 2013; Parisi et al. 2020), and NFX1−type zinc finger−containing protein 1 (RNA-binding protein that initiates the antiviral response and interacts with MAVS; Wang et al. 2019). These 107 transcripts were further evaluated using KEGG via MetaCerberus, which showed that most terms were involved in genetic information processing and signaling and cellular processes, with some terms involved in the immune response, similar to the differential gene expression analysis (Figure 6A). The timepoint model yielded 9 significant eigengene modules, with the most significant module being module 33 (ME 33; *p*=1.11e-05; Figure 6D). This module contained 111 genes and showed a shift in expression between T0 and T14, with the field population exhibiting a pattern similar to that at T14. A large proportion of genes showed high expression at the T14 timepoint and low expression at T0. Additionally, the KEGG terms are similar to those in module 35 of the genotype-region analysis, with more functional terms identified (Figure 6D).

## Discussion

Common garden experiments like the one conducted here provide an opportunity to test for patterns of variation and potential local adaptation by hosts to microbial and viral communities under natural conditions ^96–99^. *Nematostella vectensis* genotypes exhibited distinct and regionally structured associations with viruses, bacteria, and immune responses, revealing a coordinated ecological and evolutionary signature across a north–south pattern of genetic differentiation ^53,100,101^. Across all analyses, genotypes from higher latitudes maintained tighter control over their virome and microbiome through stronger immune responses, whereas lower-latitude genotypes accumulated more viruses and bacteria and showed weaker host-level regulation. Overall, these results suggest regional adaptation and indicate that host immunity, the virome, and the microbiome influence one another in shaping community composition when anemones encounter a new environment.

Our viral taxonomic analysis showed that dsDNA viruses dominated the virome both before and after mesocosm exposure (Figure 2), in contrast to previous reports of RNA-virus–dominated cnidarian viromes ^49,50^. Methodological differences likely contribute to these discrepancies, particularly our conservative taxonomic filtering, which may preferentially retain confidently identified viral dsDNA sequences while excluding highly divergent RNA viruses or host-derived transposons ^16,50,102–106^. Yet despite these methodological considerations, the ecological signal from the association with viruses was clear. After mesocosm exposure, lab-cultured animals accumulated viruses and shifted toward, but did not fully converge with, the natural population, indicating that viral acquisition in the mesocosm reflects biologically meaningful interactions rather than laboratory artifacts.

Within this broader shift, genotype- and region-specific viral signatures remained strong. Northern and southern genotypes harbored distinct viral communities, paralleling a genetic and bacterial diversity break at Cape Cod ^53,100,101^. This pattern aligns with the geographic mosaic theory of coevolution ^96,107–111^, in which selective pressures vary across environments and produce spatially structured host–microbe interactions. For *N. vectensis*, the virome and microbiome may act as selection gradients, and the host immune system represents the adaptive trait responding to these pressures. Consistent with this framework, viruses have been suggested to regulate and shape the host-associated microbiomes, making them drivers of adaptation ^9–11^. In this study, northern genotypes, which mounted a stronger immune response and maintained lower viral and microbial loads, appear better suited to the northern environment where the experiment was conducted. In contrast, southern genotypes accumulated more viruses, particularly lytic bacteriophages such as *Tequatrovirus,* and exhibited higher bacterial diversity and richness, suggesting that phage-mediated microbial regulation may partially compensate for weaker host immunity ^112–114^. Consistent with this hypothesis, studies have shown that high viral lysis can increase bacterial diversity and maintain community richness ^10,115–118^. Alternatively, the higher number of bacteriophages may simply reflect the greater bacterial diversity associated with anemones originally collected from lower-latitude locations.

Functional viral annotations reinforce an interpretation for spatially distinct host-microbe communities. After mesocosm exposure, viral proteins associated with replication, lysis, and phage structural components increased markedly, especially in southern genotypes (Figure 3; Supplemental Figures 4 and 5). These signatures are consistent with elevated lytic phage activity, a process known to enhance bacterial diversity through “kill-the-winner” dynamics ^10,116–120^. The parallel increase in bacterial richness in southern genotypes supports the hypothesis that viral activity shapes the microbiome in a genotype-specific manner, potentially influencing host fitness and environmental adaptation ^9–11,115–118^. KEGG pathways further revealed activation of innate immune signaling (e.g., RIG-I-like, NOD-like, Toll-like pathways), mirroring the expression patterns of the *N. vectensis*-specific immune gene set and underscoring the interplay between viral exposure and host immunity ^85,121–124^.

The associated bacterial community exhibited similar genotype- and region-specific patterns. While major phyla (i.e, Bacteroidota, Firmicutes, Proteobacteria, and Spirochaetota) matched previous studies ^37,53^, the composition and diversity differed sharply between northern and southern genotypes. Southern genotypes displayed significantly higher bacterial diversity and a notable absence of Firmicutes, a group commonly associated with antimicrobial production and immune modulation in cnidarians and other animals ^18,37,125–129^. The increased association of cyanobacteria across all genotypes after mesocosm exposure suggests a shared response to altered nutrient landscapes, potentially reflecting a compensatory strategy for nitrogen cycling or metabolic flexibility in a more natural environment ^130^. Yet the magnitude and direction of microbial shifts remained genotype-specific, reinforcing the idea that host genetic background shapes microbial assembly even under shared environmental conditions ^37^.

The host transcriptomic analysis indicates that genotype strongly influences host transcriptomic responses in the mesocosm. Northern genotypes exhibited a more robust immune response, with hundreds of differentially expressed genes enriched for cytokine receptors, pattern recognition receptors, antimicrobial genes, and apoptosis-related pathways, which have been linked to host defense strategies against viral infection ^85,131,132^. Southern genotypes, in contrast, showed minimal transcriptional activation, consistent with their higher viral and microbial loads. Together, these findings lead us to hypothesize that northern genotypes may rely more heavily on host-mediated immune regulation, whereas southern genotypes may depend more on microbe–virus interactions to structure their internal ecosystem, at least under these experimental conditions. Alternatively, it is possible that co-evolution of some elements in the anemone’s immune system with local bacteria made it more effective in maintaining lower levels of potential pathogens.

Together, this study supports a geographic mosaic of coevolution in which host immunity, viral dynamics, and microbial communities interact across spatially structured environments. This eco-evolutionary interplay suggests that host fitness is context-dependent, shaped by the alignment (or mismatch) between genotype-specific regulatory strategies and local microbial and viral landscapes.

## Acknowledgements

We thank the University of New Hampshire Jackson Estuarine Laboratory for providing the mesocosm and laboratory facilities used to conduct these experiments.

## Data Accessibility

The sequencing data generated in this work are available under accession PRJNA1499806 at the SRA database of NCBI. All bioinformatic workflows can be found on GitHub https://github.com/sydney-birch/Nematostella_Mesocosm_virome.

