## Supplementary Data for "Intersection of genotype and environment on the dynamics of the virome and microbiome of an estuarine cnidarian"

### I. Supplementary Figures

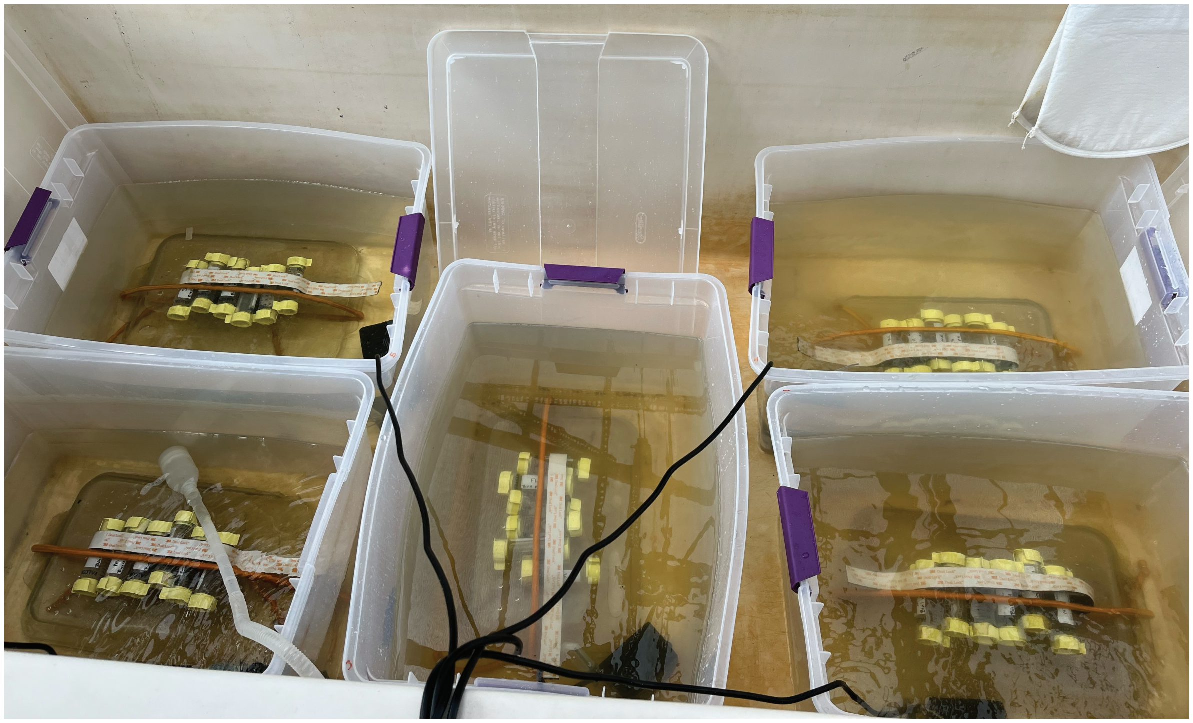

**Supplemental Figure 1. Mesocosm set up**

The five mesocosm replicates deployed at the University of New Hampshire Jackson Estuarine Laboratory (Durham, New Hampshire, USA). Each mesocosm (plastic bin) contained animal tubes of each clonal line (genotype; NS, ME, NH, MA, SC, FL) along with a water pump to create gently flowing water. Mesocosms were filled with natural estuary water that was bag-filtered to remove larger debris. Estuary water was replaced every 48 hours.

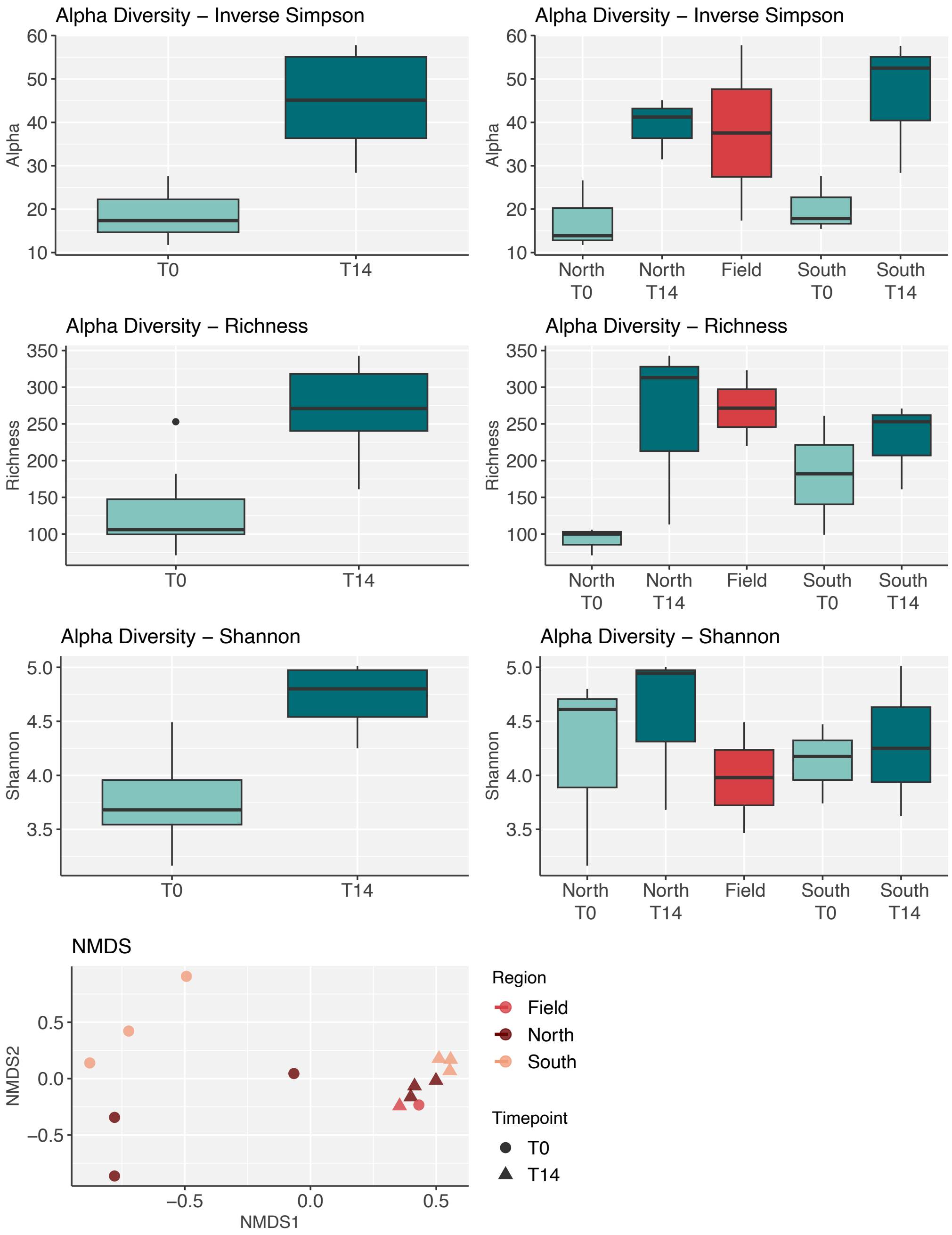


**Supplemental Figure 2. Viral Diversity Metrics**

Bar plots of three alpha diversity tests (Inverse Simpson, Species Richness, and Shannon Diversity) examining viral diversity at time point (T0 vs T14) and genotype-region by time point (North T0, North T14, etc). In the bar plots, light teal indicates T0 timepoint samples, dark teal indicates T14 timepoint samples, and red indicates the field (natural population) samples. Additionally, we examined beta diversity using an NMDS, where the time point is indicated by the shape of the symbol and the genotype region is indicated by color in the plot.

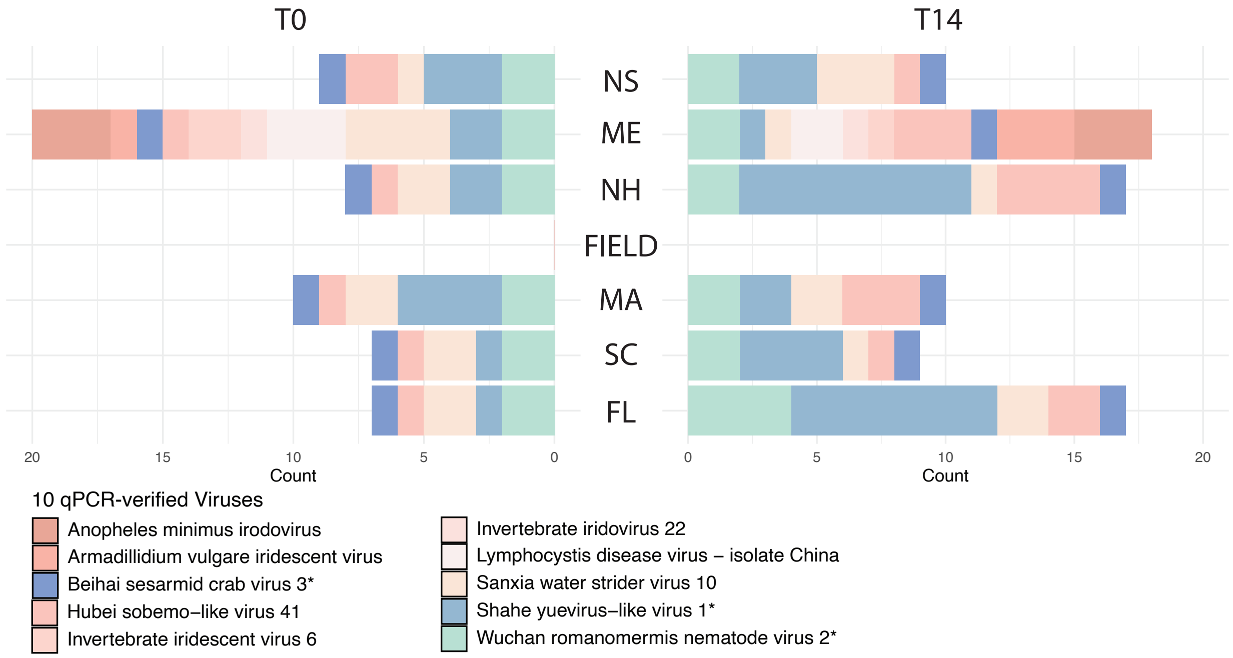


**Supplemental Figure 3. Artemia Viruses identified from Lewandowska et al, 2020**

Stacked bar plot depicting counts of the 10 qPCR-verified core viruses from Lewandowska et al, 2020, identified in our dataset across genotypes and time points. Viruses labeled in blue-green colors with asterisks next to their names are Artemia viruses, while those labeled in pink indicate other viruses.

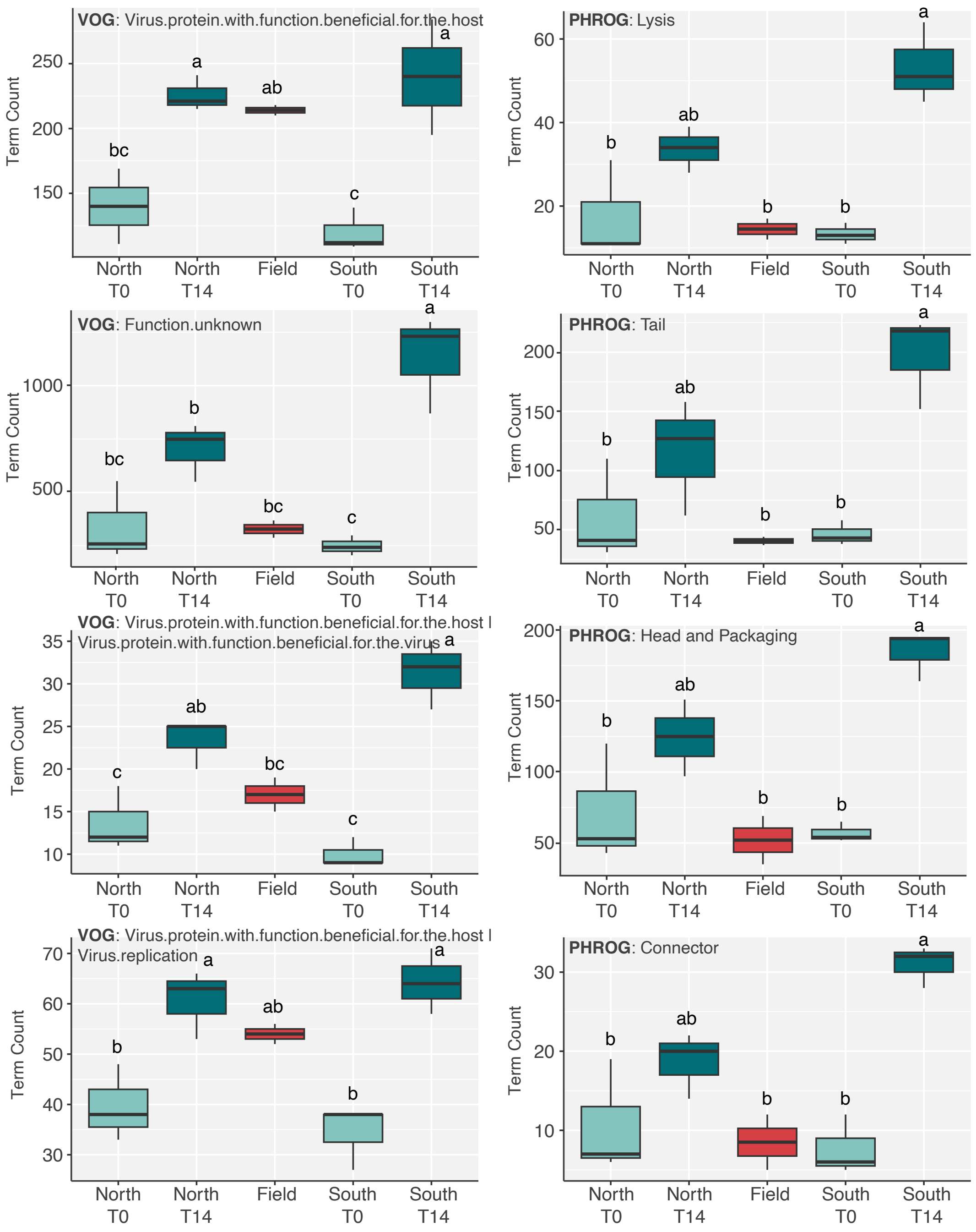


**Supplemental Figure 4. Statistics of viral functional terms from PHROG and VOG databases**

Box plots were generated for terms of interest from PHROG and VOG databases. We examined genotype-region by time point (North T0, North T14, etc.) using ANOVA followed by Tukey's HSD to assess significance. In the bar plots, light teal indicates T0 timepoint samples, dark teal indicates T14 timepoint samples, and red indicates the field (natural population) samples. Each bar plot contains the term database in bold, followed by the term as the title.

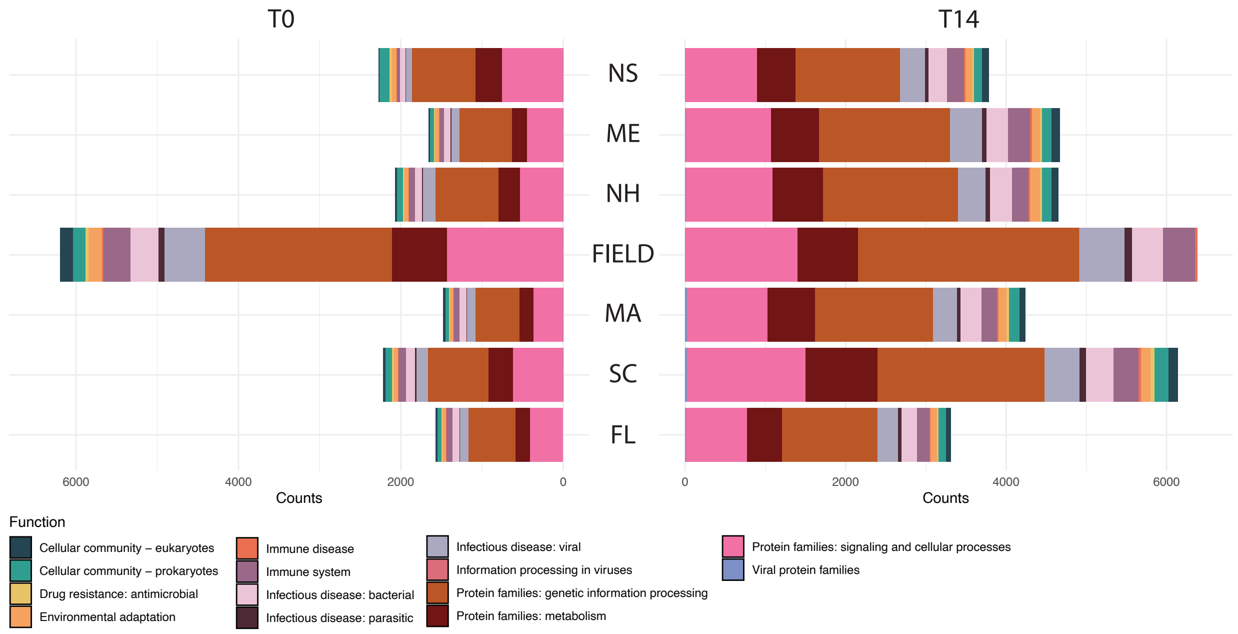


**Supplemental Figure 5. KEGG Functional Term Counts for Viruses**

Mirrored bar plot of select KEGG functional term counts for viruses across genotypes and time points. ANOVAs were run on select terms interrogating time point: genetic information processing (p=0.0002), infectious disease: viral (p=6.1e-05) and bacterial (p=6.82e-05), immune system (p=0.00023), and environmental adaptation (p=2.62e-05).

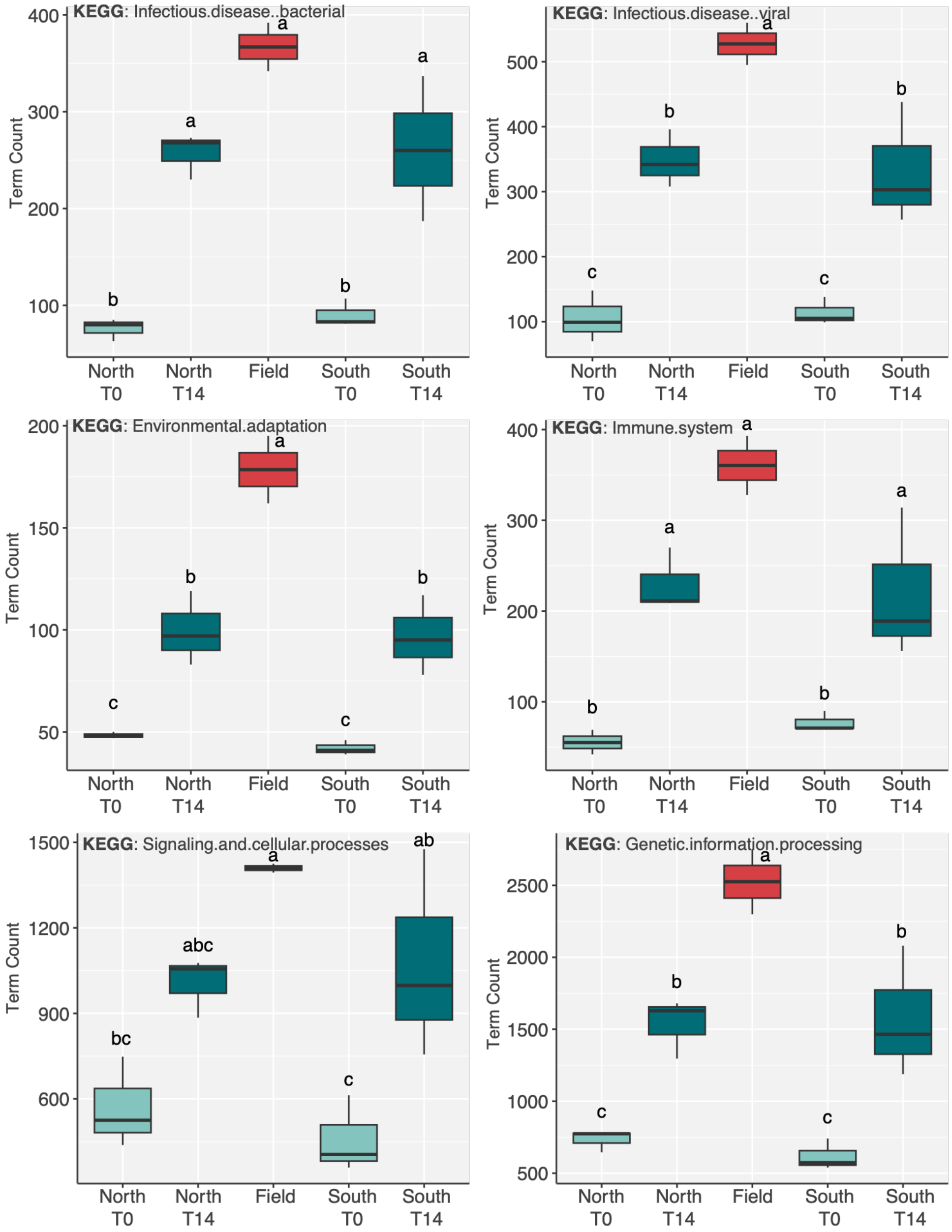


**Supplemental Figure 6. Statistics of viral functional terms from KEGG database**

Box plots were generated for terms of interest from the KEGG database. We examined genotype-region by time point (North T0, North T14, etc.) using ANOVA followed by Tukey's HSD to assess significance. In the bar plots, light teal indicates T0 timepoint samples, dark teal indicates T14 timepoint samples, and red indicates the field (natural population) samples. Each bar plot contains the term database in bold, followed by the term as the title.

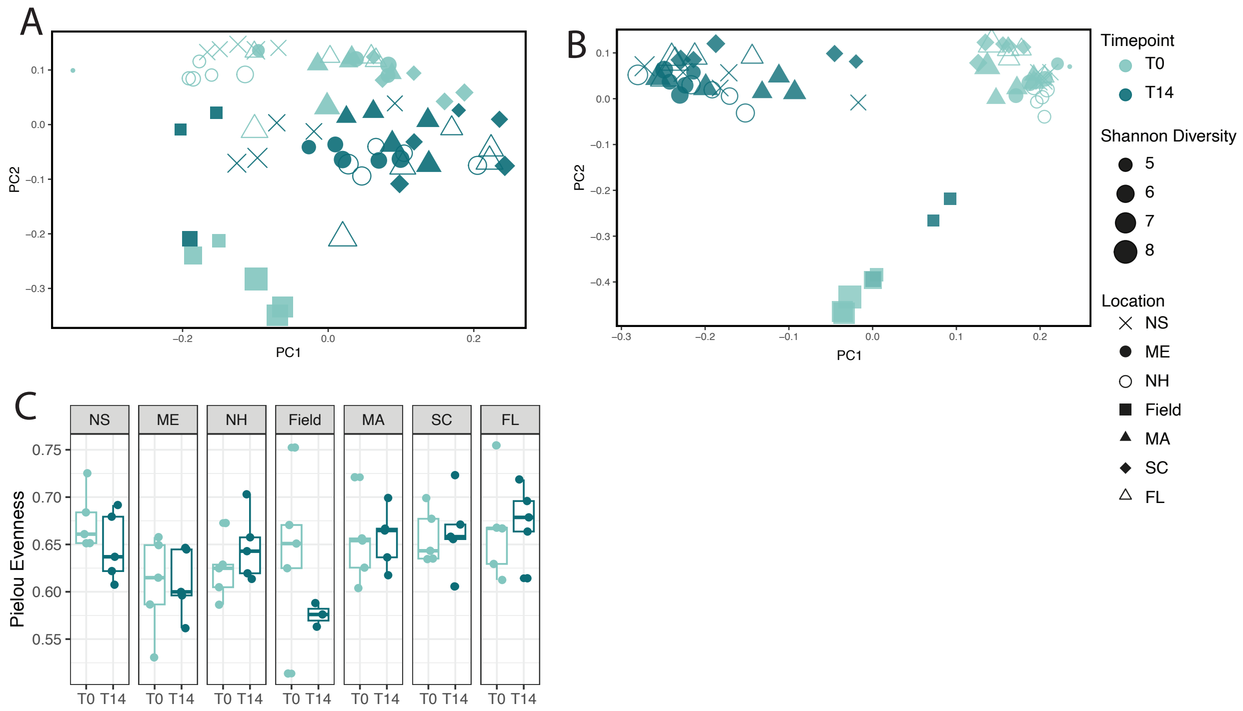


**Supplemental Figure 7. Additional Bacterial Diversity Analyses**

Additional bacterial diversity analyses, generated in QIIME2. (**A**) Unweighted unifrac and (**B**) Jaccard distance PCAs examining bacterial beta diversity. Symbol shape indicates the genotype location, color indicates time point (light teal: T0, dark teal: T14), and size of the symbol indicates the Shannon diversity index. (**C**) Box plot of Pielou's Evenness index assessing bacterial evenness across genotype locations and time points. Light teal indicates T0 timepoint samples, dark teal indicates T14 timepoint samples.

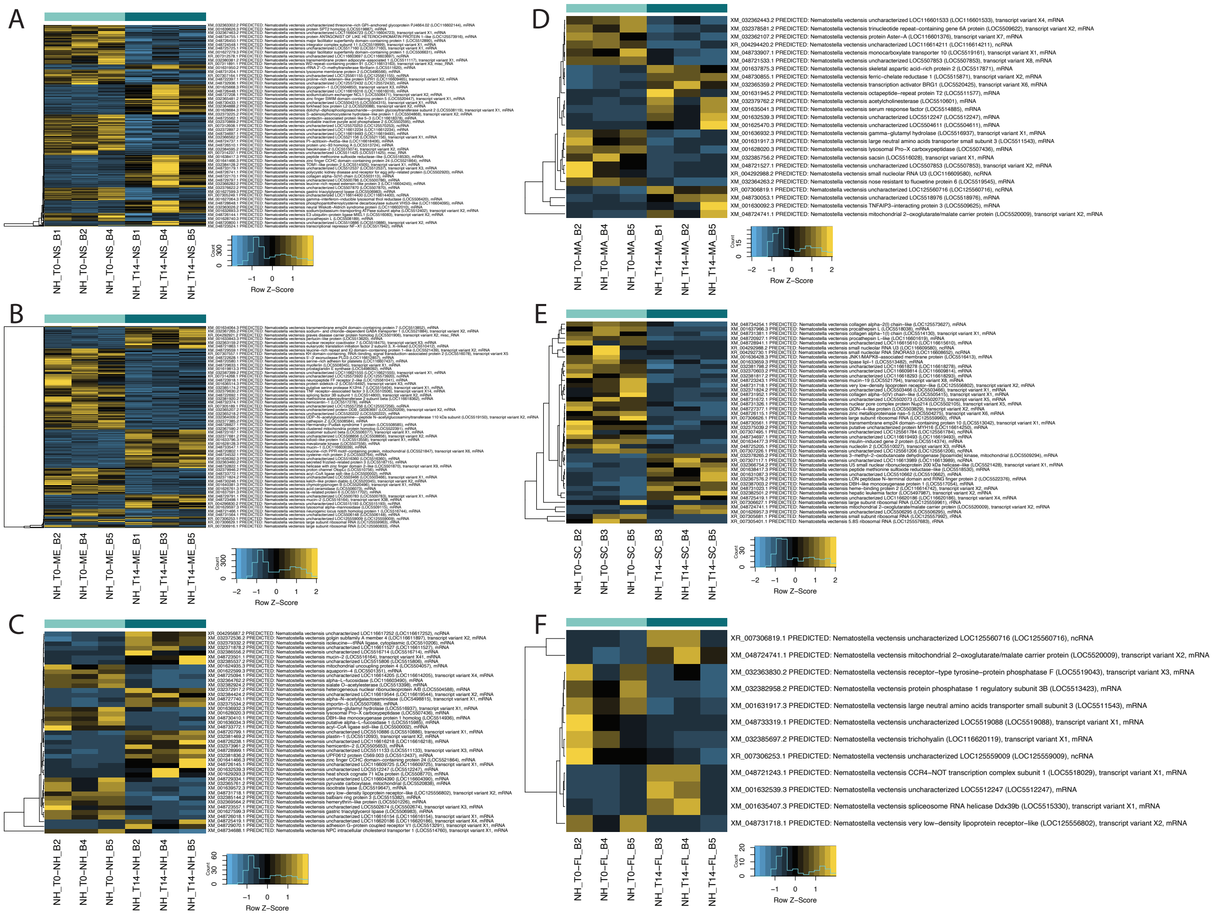


**Supplemental Figure 8. Heatmaps of Host (*N. vectensis*) Differentially Expressed Genes**

Differentially expressed genes were identified using edgeR by comparing expression between time points for each genotype: (**A**) Nova Scotia (469 DEGs), (**B**) Maine (377 DEGs), (**C**) New Hampshire (43 DEGs), (**D**) Massachusetts (25 DEGs), (**E**) South Carolina (43 DEGs), and (**F**) Florida (12 DEGs). Each row corresponds to a single transcript, labeled with the annotated header from the UK transcriptome. Yellow indicates high expression, and light blue indicates low expression. The teal bars above the heatmap indicate time points where light teal indicates T0 samples and dark teal indicates T14 samples.

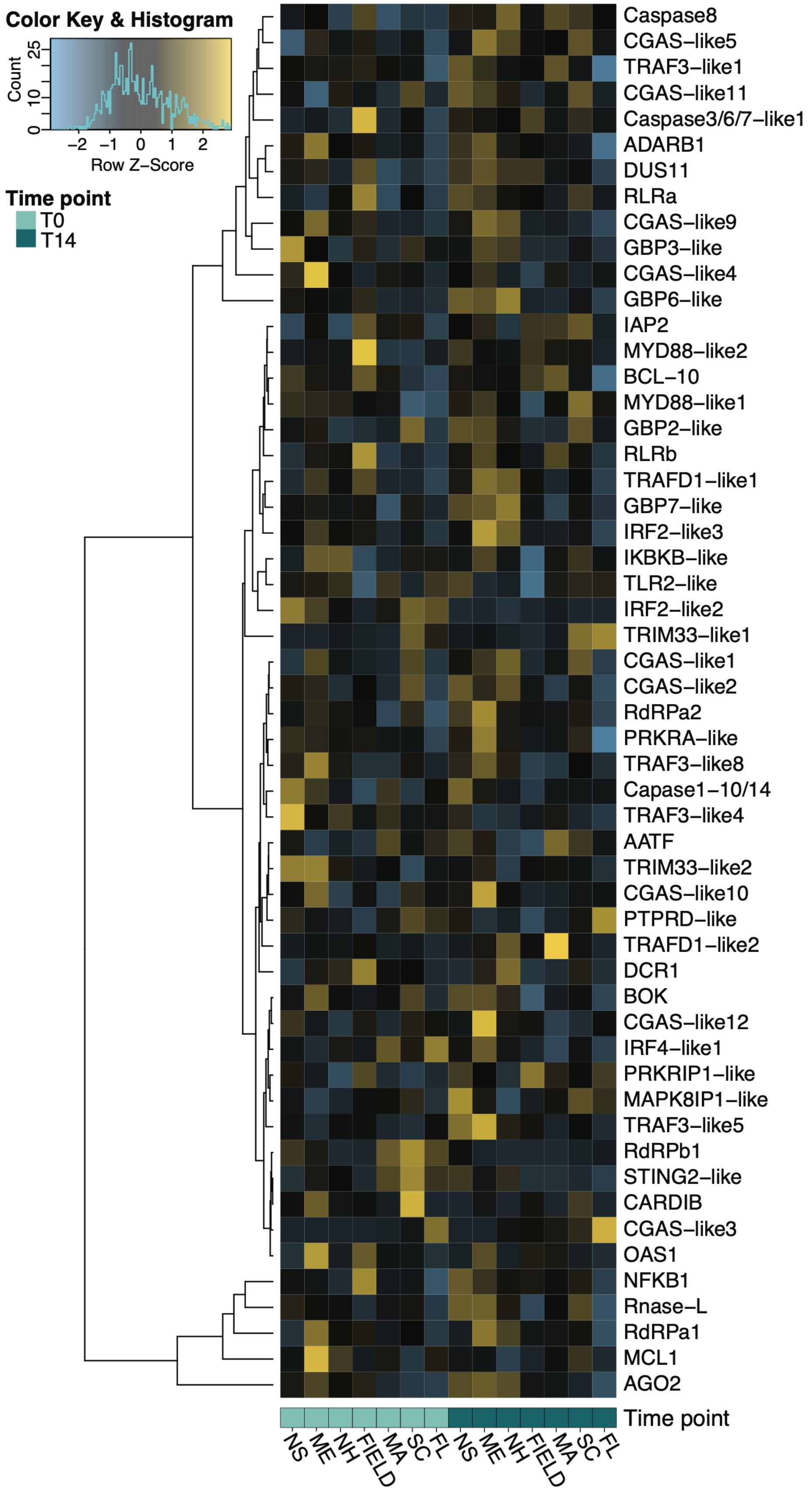
Color Key & Histogram
Count
0
10
25
−2 −1 0 1 2
Row Z−Score
Time point
T0
T14
Caspase8
CGAS−like5
TRAF3−like1
CGAS−like11
Caspase3/6/7−like1
ADARB1
DUS11
RLRa
CGAS−like9
GBP3−like
CGAS−like4
GBP6−like
IAP2
MYD88−like2
BCL−10
MYD88−like1
GBP2−like
RLRb
TRAFD1−like1
GBP7−like
IRF2−like3
IKBKB−like
TLR2−like
IRF2−like2
TRIM33−like1
CGAS−like1
CGAS−like2
RdRPa2
PRKRA−like
TRAF3−like8
Capase1−10/14
TRAF3−like4
AATF
TRIM33−like2
CGAS−like10
PTPRD−like
TRAFD1−like2
DCR1
BOK
CGAS−like12
IRF4−like1
PRKRIP1−like
MAPK8IP1−like
TRAF3−like5
RdRPb1
STING2−like
CARDIB
CGAS−like3
OAS1
NFKB1
Rnase−L
RdRPa1
MCL1
AGO2
Time point
NS ME NH FIELD MA SC FL NS ME NH FIELD MA SC FL

**Supplemental Figure 9. Host Expression of previously curated *N. vectensis* 56 immune-related genes**

Heatmap of immune-related gene expression previously curated in *N. vectensis* (Sharoni et al. 2026) in mesocosm and field-collected *N. vectensis*. Yellow indicates high expression, and light blue indicates low expression. The teal bars below the heatmap indicate time points where light teal indicates T0 samples and dark teal indicates T14 samples.

### II. Supplementary Tables

| Location of Origin | GPS Coordinates |
|---|---|
| Crescent Beach, Nova Scotia (NS) | 45.154713, -64.371314 |
| Saco River, Maine (ME) | 43.542944, –70.344694 |
| Wallis Sands, New Hampshire (NH) | 43.028656, –70.730775 |
| Sippewissett, Massachusetts (MA) | 41.589389, –70.637972 |
| Georgetown, South Carolina (SC) | 33.330722, -79.201167 |
| St. Augustine, Florida (FL) | 29.726417, -81.257722 |

**Supplemental Table 1. Coordinates of collection sites for clonal lines**

The *Nematostella vectensis* clonal lines used in this study were originally collected from these locations. These lines were maintained under laboratory conditions prior to being used in the mesocosm field study.
